# *In-Silico* Characterization of *TP53* Splice Mutations in Somatic and Germline Tumours

**DOI:** 10.1101/2025.05.27.656522

**Authors:** Apeksha Arun Bhandarkar, Noah Ethan Kelly-Foleni, Debina Sarkar, Aaron Jeffs, Tania Slatter, Anthony Braithwaite, Sunali Mehta

**Affiliations:** Department of Pathology, University of Otago, Dunedin, New Zealand; Maurice Wilkins Centre for Biodiscovery, University of Auckland, Auckland, New Zealand; Medical Laboratory Science, University of Otago, Dunedin, New Zealand

**Author notes:** Corresponding author Dr. Sunali Mehta, Department of Pathology, Dunedin School of Medicine, University of Otago, 56 Hanover Street, Dunedin - 9016, New Zealand.

**Keywords:** p53, mutation, splice, isoforms

## Abstract

*TP53* undergoes alternative splicing to produce multiple mRNA transcripts and protein isoforms, yet the effects of splice site mutations on isoform regulation, tumor-biology, and clinical outcome remain unclear. Analysis of 23,017 *TP53* variants, including 18,562 somatic mutations (pan-cancer datasets – cBioPortal) and 4,455 germline variants (IARC database), identified recurrent donor (X32, X125, X224, X261, X331) and acceptor (X33, X126, X187, X225, X307, X332) splice site mutations. Germline variants showed nucleotide-specific transition biases. Most splice site mutations were associated with reduced *TP53* mRNA expression; however, X32, X33, X126, and X261 maintained or elevated transcript levels. Splice mutations were associated with distinct transcriptional subsets marked by altered p53 target gene expression, elevated tumor mutation burden, increased genomic instability, and significantly reduced disease-free survival compared to missense mutations, with X126 and X331 being associated with poorest outcomes. These findings emphasize the clinical impact of *TP53* splice site mutations and the need for functional classification.

## Introduction

Splicing defects play a pivotal role in cancer by disrupting the expression of key genes associated with various ‘hallmarks of cancer’.^1–3^ These defects often arise from mutations in splicing factors e.g., SF3B1, U2AF1 or cis-regulatory elements, leading to dysfunctional proteins, oncogenic isoforms, or nonfunctional transcripts. [reviewed in^3^] Splice site mutations disrupt normal splicing, causing intron retention and aberrant transcript formation. Canonical splice site mutations at GT-AG dinucleotides further exacerbate splicing defects, leading to exon skipping, intron retention, or activation of cryptic splice sites. Several pan-cancer studies have highlighted synonymous mutations enriched in *TP53* at critical splice regions, including the 5’ end of exon 6 and the 3’ end of exons 4 and 9.^4–6^

Analysis of large-scale cancer genomics datasets using open-access platforms like cBioPortal and the IARC TP53 Database shows that ∼7% of reported *TP53* mutations occur at canonical splice sites or splice regions (Figure. 1A and 1B; Table 1). These mutations are significant due to the complexity of the *TP53* locus, which encodes multiple transcripts (Figure 1C). Splice mutations can affect transcription from an internal promoter in intron 4 (P2), producing Δ133p53 isoforms, and the generation of Δ40p53 transcripts, regulated by G-quadruplex structures in intron 3 and splicing of intron 2. They can also lead to alternative 3′ ends between exon 9 and exons 9β/10, resulting in *TP53α/*β/γ variants (Figure 1C).^1,2,7^ Beyond protein-coding transcripts, noncoding isoforms like p53Ψ, which can promote metastatic-like behaviour, have been reported(Figure 1C).^8^ Detailed insights into the frequency and effects of splice mutations on p53 function, isoform expression, tumour biology, and patient outcome remain limited, highlighting the need for further studies to unravel the molecular and clinical significance of *TP53* splice site mutations.

**Figure 1:**
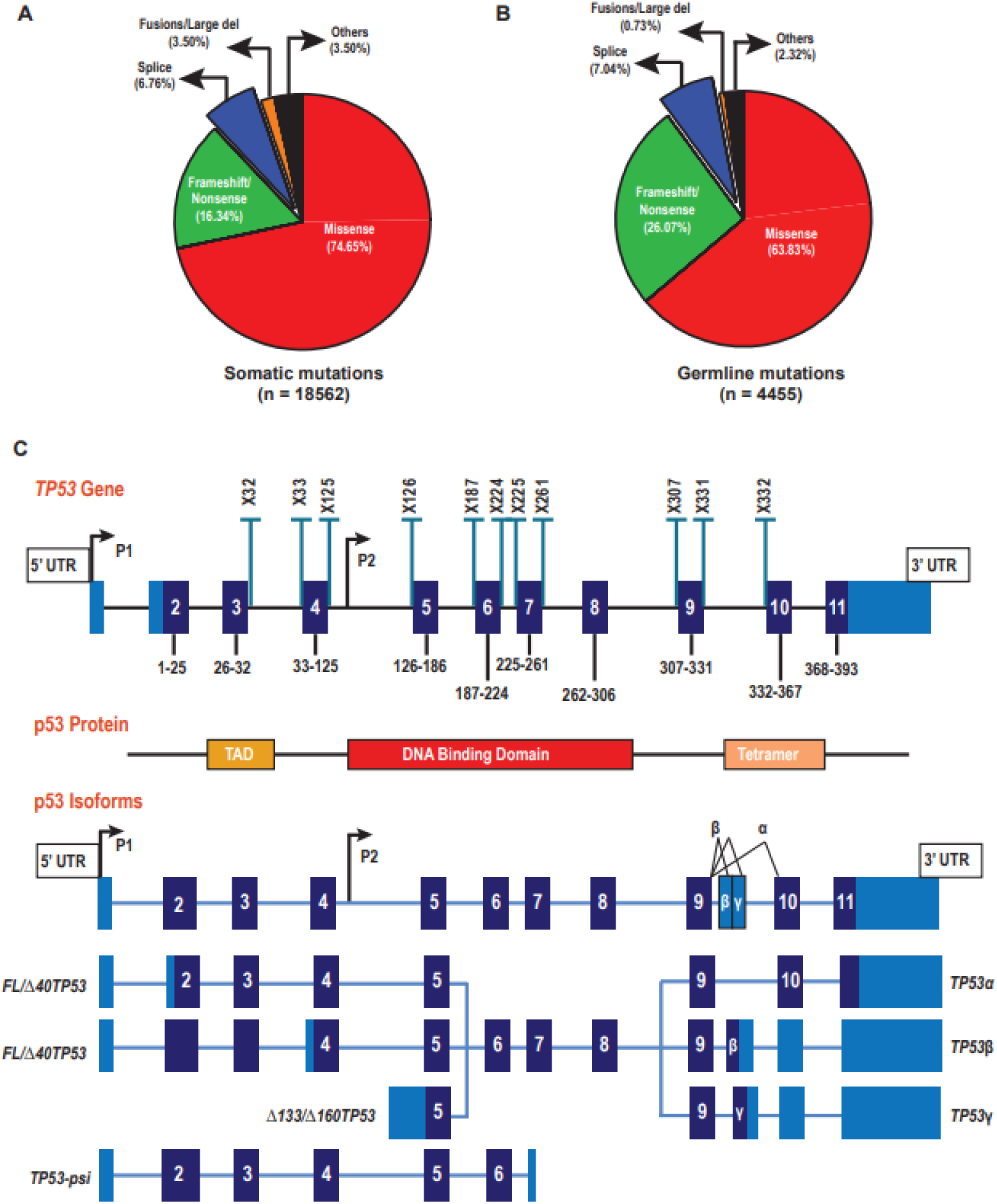
T***P***53 **mutation spectrum and location of *TP53* splice mutations in relation to p53 isoforms. A-B.** The pie chart represents the distribution of mutation types (Missense, Frameshift/nonsense, Splice, Fusions/Large Del and Others - In-frame, translation start site and nonstop mutations) reported in **A.** Somatic (TCGA Pan-Atlas Cancer Dataset, MSK-MetTropism Metastatic Dataset and Tumour Mutation Burden (TMB) and Immunotherapy Dataset) and **B.** Germline (IARC database). **C.** Schematic representing the structure of the *TP53* gene, protein and known isoforms along with the location of the splice mutations reported in the somatic and germline databases. **Top Panel:** *TP53* gene (light blue – 5’UTR and 3’UTR, dark blue – coding exons) with location of splice mutations (Blue bars). P1 and P2 are indicate promoters 1 and 2 of *TP53*. **Middle panel:** p53 protein domains. TAD: transactivation domain. **Bottom Panel:** *TP53* transcripts encoding *FLTP53α/β/γ*, *Δ40TP53α/β/γ*, *Δ133/Δ160TP53α/β/γ* protein isoforms (light blue – 5’UTR and 3’UTR, dark blue – coding exons).

This study presents for the first time a comprehensive data mining analysis of *TP53* splice site mutations in somatic and germline cancers, using data from multiple databases. We show that one subset of splice site mutations leads to loss of p53 function, and is associated with higher tumour mutation burden, increased genomic instability, and poorer survival outcomes. Another subset also results in selective loss of p53 function but does not increase genomic instability yet is still linked to poor survival. Notably, we observe distinct molecular and clinical impacts between donor and acceptor splice site mutations flanking specific introns, underscoring the complexity and clinical relevance of these alterations.

## Methods

### Retrieval of somatic *TP53* mutations reported in the cBioPortal Database

Integrative analysis of reported somatic *TP53* mutations across all cancer types was performed using cBioportal^9–11^ – a publicly available database for tumour genetics and transcriptomics (accessed and downloaded in September, 2024). This database allows large scale statistical analysis and graphical viewing of tumour changes at mRNA and protein level. We based our study on data generated by The Cancer Genome Atlas (TCGA) Pan-Atlas Cancer Dataset^12^, Memorial Sloan-Kettering (MSK)-MetTropism Metastatic Dataset^13^ and Tumour Mutation Burden (TMB) and Immunotherapy Dataset (TMBID)^14^. A total of 11378 samples were derived from the TCGA dataset, out of which 7128 were genetically wild type (WT) and 4250 carried a *TP53* gene mutation (∼37%). Similarly, a total of 27696 samples were derived from the MSK-MetTropism dataset, out of which 13325 were genetically WT and 13471 had a *TP53* gene mutation (∼49%). Lastly, form the TMBID dataset 1764 sample details were derived, with 923 as genetically WT and 841 having *TP53* gene mutations (∼48%). Each of these datasets had few patient IDs in duplicates associated with different sample IDs (TCGA - 411 duplicates, MSK-MetTropism – 1021 duplicates, TMBID – 103 duplicates). Each sample ID was considered as an individual datapoint.

### Retrieval of germline *TP53* mutations reported in the International Agency for Research on Cancer (IARC) *TP53* Database

The reported germline *TP53* mutations was downloaded from IARC database^15^ (accessed and downloaded in October 2024) along with their characteristics and family history. The IARC database is a publicly available database for the analysis, interpretation of biological and clinical impacts of *TP53* mutations on human cancers. A total of 4455 mutations were retrieved from the database under germline variants, with multiple variants reported under the same family ID.

### *TP53* mutation frequency, and their molecular and clinical characteristics

We categorised the *TP53* mutations from somatic and germline datasets as missense, frameshift/nonsense, splice, fusions/large deletions and other mutations (in-frame, translation start site and nonstop mutations). The most frequently occurring mutations in *TP53* were considered as hotspot mutations namely R175, G245, R248, R273, R282 and Y220^16^ and are included as part of the missense mutations. For our analysis, the data from the three somatic datasets (TCGA, MSK-MetTropism and TMBID) from cBioPortal^9–11^ were combined under somatic category. For the germline data, single nucleotide changes occurring at the first or last nucleotide of exons were considered as missense mutations, though labelled as splice effect (85 mutations).

The main splice mutations (X32, X33, X125, X126, X187, X224, X225, X261, X307, X331, X332) were selected based on a frequency threshold of >=3% of the total splice mutations in each dataset. To assess the frequency and distribution of *TP53* splice site mutations across tumour types, including bladder, breast, colorectal, esophagogastric, head and neck, melanoma, non-small cell lung cancer, and renal cell carcinoma, we analysed the types of genetic alterations at each splice site position in both somatic and germline datasets. These included single nucleotide polymorphisms (SNPs), dinucleotide polymorphisms (DNPs), deletions (del), and insertions (ins). However, for splice sites, only SNPs occurring at these sites were included in the analysis, and sub-categorised as transversions (C>A, C>G, A>T, A>C) and transitions (C>T, A>G), to compare patterns between the somatic and germline datasets.

The molecular characteristics analysed included mutation count and fraction of the genome altered (FGA), which were available only in the somatic datasets namely TCGA and MSK-MetTropism. Mutation count refers to the total number of mutations present in a tumour sample, while FGA represents the percentage of the genome affected by changes such as copy number variations, structural rearrangements, insertions, deletions, and single nucleotide variants. Together, these metrics provide an overall measure of genomic instability. Study IDs and sample IDs for each *TP53* mutation category, including splice site mutations, were retrieved from the raw data and used for statistical analysis of mutation count and genomic alteration. The TCGA and MSK-MetTropism datasets were used for this study, focusing on only the primary sample types and excluding the sample types which are labelled as metastasised. Further the clinical characteristics analysed included the disease-free survival (available only in the TCGA dataset). Disease-free survival is the period from the date of diagnosis until the date of first recurrence, loco-regional or systemic. This data was retrieved from TCGA dataset for all *TP53* mutation categories and analysed.

mRNA and protein expression levels were assessed in primary tumour samples using data from The Cancer Genome Atlas (TCGA). Expression distributions of *TP53* splice site mutations were compared to those of wild-type *TP53* using the density function in R (version 4.4.3^17^), a non-parametric method for estimating probability density, allowing visualization of expression patterns without assuming an underlying distribution.

To investigate the impact of *TP53* splice site mutations on p53 signalling, a reference list of 663 genes known to be *TP53* targets or associated with *TP53* expression was compiled (Table S1). This list was curated using multiple resources available through the Enrichr database^18–20^, including the ARCHS4 RNA-seq gene-gene co-expression matrix^21^, Enrichr gene-gene co-occurrence matrix^18–20^, Tagger literature gene-gene co-mentions matrix^22^, and GeneRIF literature gene-gene co-mentions matrix^23^. mRNA expression data for these genes were extracted from the TCGA Pan-Cancer dataset for tumour samples harbouring *TP53* splice site mutations. Expression values were log-transformed and mean-centred to normalize across samples. Hierarchical clustering (Euclidean distance and complete linkage) was performed to group samples based on similarity in gene expression profiles. Clustering was applied to both rows (genes) and columns (samples), allowing the identification of distinct expression patterns and potential subgroups among *TP53* splice mutant tumours. Heatmaps were generated to visualize co-expression patterns and to explore the association between *TP53* splice site mutations and transcriptional dysregulation of p53-associated genes. R (version 4.4.3^17^) - was used to normalise the data, perform hierarchical clustering and generate a heatmap.

### Statistical Analysis

Fisher’s exact test was applied to assess whether there was a significant difference in the nucleotide change between somatic and germline mutations, as well as individual base changes contributing to specific splice mutations. All categories of *TP53* mutations, including splice mutations, were statistically analysed with respect to mutation count and FGA. The Kruskal-Wallis test was applied to assess whether significant differences (p-value < 0.05) existed between the different *TP53* mutation groups. To evaluate differences in disease-free survival, the log-rank test was performed. For mRNA and protein expression levels, the Kolmogorov-Smirnov (KS) test was used to determine whether they significantly differed between tumours with *TP53* mutations and those without. Fisher’s exact test was employed to assess the significance of splice mutation frequency, while the Chi-square test for trends was used to evaluate the distribution of splice mutations by tumour type across the two clusters identified in the heatmap. Statistical analyses and graphical representations were conducted using GraphPad Prism (version 10.4.2), R (version 4.4.3^17^), and Adobe Illustrator (version 29.5.1 (64 bit)).

## Results

### Frequency and characteristics of reported *TP53* splice mutations in somatic and germline tumours

Reported *TP53* splice site mutations are present at both donor and acceptor sites flanking introns 3 (X32– X33), 4 (X125–X126), 6 (X224–X225), and 9 (X331–X332); at the acceptor sites flanking introns 5 (X187) and 8 (X307); and at the donor site flanking intron 7 (X261). These mutations occur in both somatic and germline tumours but at varying frequencies (Figure 1C and Figure 2A and 2B). A significant difference in mutation frequency between somatic and germline datasets is observed (Chi-squared test, p < 0.0001), with X33, X187, X224, X225, X331 and X332 more frequently found in germline tumours, whereas X32, X126, X261 and X307 are more common in somatic tumours (Figure 2A).

**Figure 2:**
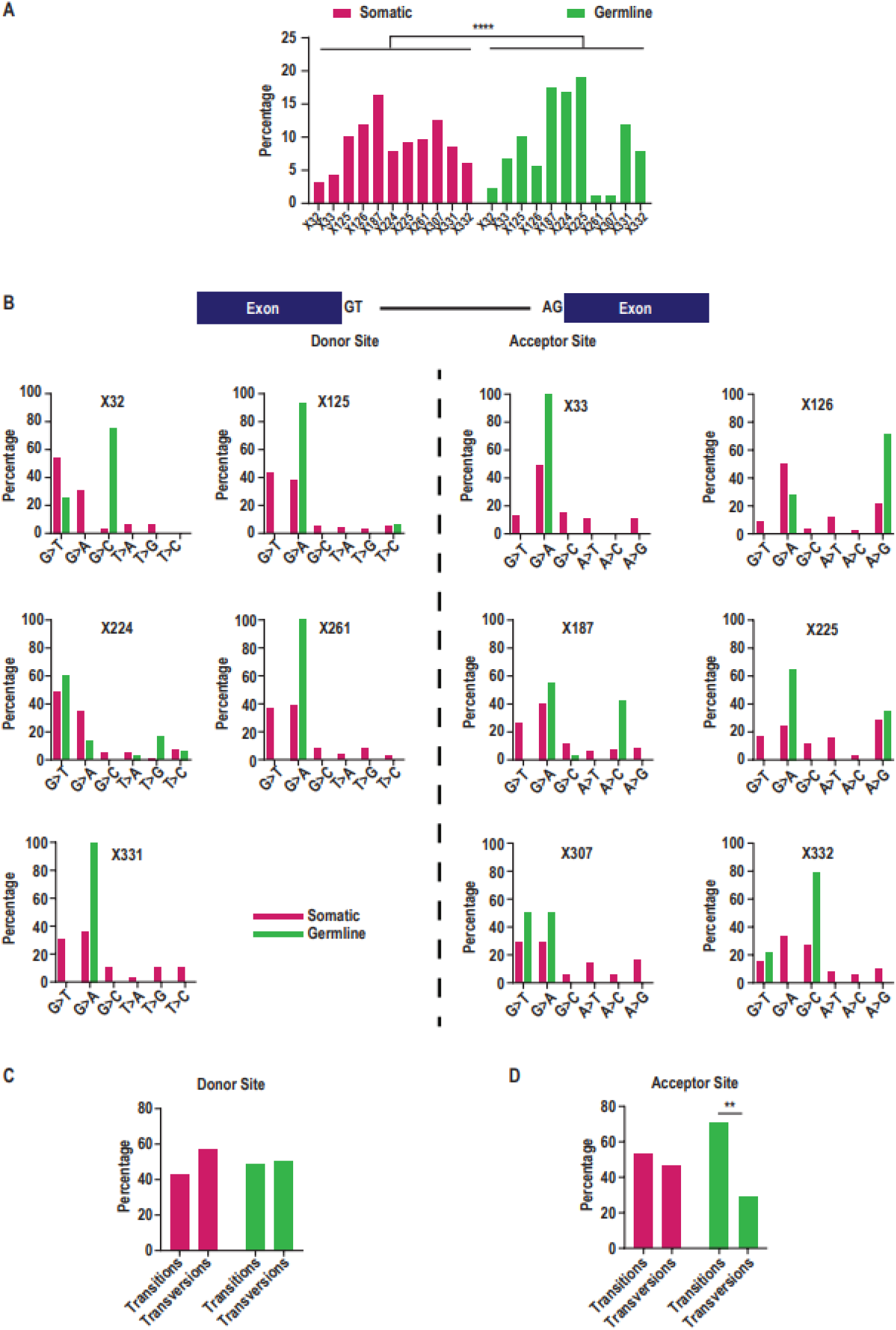
T***P***53 **splice mutations and their characteristics from somatic and germline databases. A.** Bar chart showing the percentage distribution of individual *TP53* splice mutations across Somatic (TCGA Pan-Atlas Cancer Dataset, MSK-MetTropism Metastatic Dataset and Tumour Mutation Burden (TMB) and Immunotherapy Dataset) and the germline (IARC database). Significance was determined using (Chi- square test) and p <0.05 is considered significant. **B.** Schematic showing the bases that constitute donor and acceptor splice sites flanking exons. Bar graphs depicting the percentage of base substitution types— G>A/T/C and T>A/G/C at donor sites, and A>T/C/G and G>T/A/C at acceptor sites—for individual *TP53* splice site mutations. C-D. Bar graph showing the percentage of base substitutions resulting in transitions or transversions in C. Donor sites and D. Acceptor sites. Donor site mutations include X32 (somatic: n=40, germline: n=4), X125 (somatic: n=124, germline: n=18), X224 (somatic: n=97, germline: n=30), X261 (somatic: n=120, germline: n=2), and X331 (somatic: n=104, germline: n=21), while acceptor site mutations include X33 (somatic: n=54, germline: n=12), X126 (somatic: n=146, germline: n=10), X187 (somatic: n=201, germline: n=31), X225 (somatic: n=114, germline: n=34), X307 (somatic: n=153, germline: n=2), and X332 (somatic: n=75, germline: n=14). Significance was determined using (Fisher’s Exact test) and * p <0.05 - significant.

The canonical nucleotide sequence at donor splice sites is typically GT/CA, while acceptor sites usually consist of AG/TC^24,25^ (Figure 2B, top panel). Nucleotide substitutions such as G>A/T/C and T>A/G/C are commonly observed at donor sites, whereas A>T/C/G and G>T/A/C mutations are reported at acceptor sites, each with varying frequencies in germline and somatic datasets. Among donor site mutations, the nucleotide immediately adjacent to the exon (“G”) is more frequently mutated, with less than 10% of mutations occurring at the second nucleotide (“T”). Furthermore, the frequency and distribution of G>T, G>A, and G>C substitutions at donor sites differ between somatic and germline tumours. G>T mutations resulting in X32 and X224 are observed in both somatic and germline tumours, whereas the same mutation at X125, X261, and X331 is reported only in somatic tumours (Figure 2B, donor sites). In contrast, G>A mutations at X125, X261, and X331 are seen in both datasets, but G>A mutations leading to X32 are exclusive to somatic tumours (Figure 2B, donor sites). G>A mutations resulting in X224 are found in both somatic and germline tumours, although the frequency is lower in the germline (Figure 2B, donor sites). G>C mutations are the least frequently observed substitution at donor sites, with the exception of a G>C mutation at X32 reported in germline tumours (Figure 2B, donor sites).

At acceptor sites, a similar pattern is seen, the nucleotide closest to the exon (“G”) is more frequently mutated than the second position (“A”), with the latter accounting for less than 10% of reported mutations (Figure 2B, acceptor sites). Notable exceptions include A>G mutations at X126 and X225, and an A>C mutation at X187 (Figure 2B, acceptor sites). Interestingly, the A>G change at X126 is more prevalent in germline than somatic tumours, whereas this trend is not observed for X225 (Figure 2B, acceptor sites). The A>C mutation at X187, similar to X126, is more common in germline tumours (Figure 2B, acceptor sites). Differences in the frequency of G>T, G>A, and G>C changes at acceptor sites are also apparent (Figure 2B, acceptor sites). For example, G>T mutations leading to X33, X126, X187, and X225 are observed exclusively in somatic tumours, while G>T mutations at X307 and X332 occur in both somatic and germline datasets. Among the three substitution types, G>A is the most reported, occurring at most acceptor sites in both datasets, except for X332, where it is absent in germline (Figure 2B, acceptor sites). In contrast, G>C is the most frequently observed alteration in germline tumours at the X332 acceptor site (Figure 2B, acceptor sites).

In addition to these site-specific patterns, broader trends are also evident. In the germline dataset, no significant bias is observed at donor splice sites (Figure 2C); however, transitions are significantly more frequent than transversions at acceptor sites (Fisher’s exact test p < 0.0011, Figure 2D), indicating a site-specific pattern. In somatic datasets, donor sites show a trend toward a higher frequency of transversions over transitions, although this trend does not reach statistical significance (Figure 2C and 2D).

These findings underscore distinct mutational patterns at *TP53* splice sites, highlighting not only positional and nucleotide-specific biases, but also key differences in mutation spectra between somatic and germline tumours, which may reflect underlying biological mechanisms or selective pressures in tumour development.

### Impact of *TP53* splice mutations on *TP53* RNA and protein expression

To investigate the effect of splice mutations on tumour biology, we first asked whether the presence of splice mutations alters *TP53* RNA and protein expression. We analysed data from the pan-cancer TCGA dataset. As previously reported^26^, *TP53* RNA and protein levels were elevated in tumours with missense *TP53* mutations but reduced in those with frameshift or nonsense mutations compared to tumours with wild-type (WT) *TP53* alleles (Figure 3A and Figure S1A). Notably, tumours with *TP53* splice mutations also showed significantly lower mRNA levels than WT tumours overall (Figure. 3A). However, unlike frameshift or nonsense mutations, a subset of splice-mutant tumours exhibited mRNA levels comparable to WT tumours, suggesting heterogeneity in their impact on *TP53* expression.

**Figure 3:**
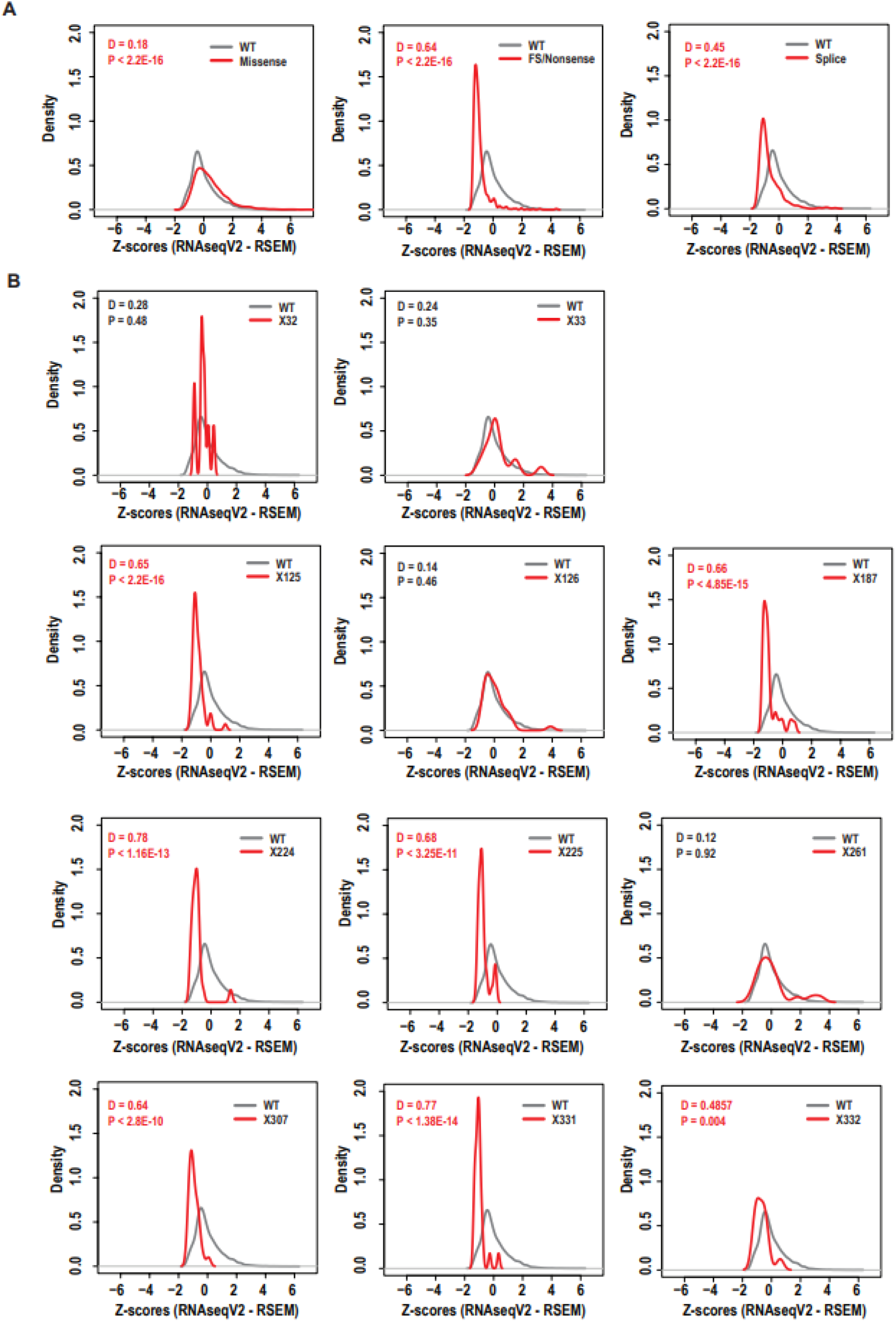
I**m**pact **of *TP53* splice mutations on *TP53* mRNA Expression. A-B.** Density plot of *TP53* mRNA expression from the RNA-sequencing of the TCGA pan-cancer dataset. **A.** Tumours with WT-*TP53* (n=6084, grey) versus different mutation types (red), left panel: Missense (n=2268), middle panel: frameshift/nonsense (n=954) and right panel: splice (n=300). **B.** Tumours with WT-*TP53* ( n=6084, grey grey) versus *TP53* splice mutations (red) as follows: X32 (n=9), X33 (n=15), X125 (n=54), X126 (n=35), X187 (n=38), X224 (n=25), X225 (n=27), X261 (n=20), X307 (n=27), X331 (n=27) and X332 (n=13). Kolmogorov-Smirnov (KS) test was used to determine significance. p < 0.05 - significant.

To explore this further, we examined individual *TP53* splice mutations and found considerable variability in *TP53* mRNA and protein expression across these tumour samples. Most splice mutations, including X125, X187, X224, X225, X307 and X331, were associated with lower mRNA expression compared to WT (Figure. 3B). However, apart from X125 and X225, which also had reduced protein levels, there were no significant changes in p53 protein levels for these mutations (Figure S1B).

In contrast, tumours with splice mutations X32, X33, X126, and X261 showed mRNA expression like WT. Consistent with this, X126 and X261 mutations were also associated with increased *TP53* protein levels (Figure S1B). Interestingly, despite significantly reduced mRNA levels, the X332 mutation was associated with significantly elevated *TP53* protein levels (Figure S1B).

These findings highlight the diverse effects of *TP53* splice mutations on p53 mRNA and protein expression and suggest that individual splice variants may differentially influence p53 levels through mechanisms such as altered mRNA stability or translation efficiency.

### Differential impact of *TP53* Splice mutations on p53 signalling

To investigate the impact of individual *TP53* splice site mutations on p53 signalling, we analysed the mRNA expression profiles of 663 genes known to be direct transcriptional targets of *TP53* or associated with its regulatory network. This analysis was conducted using hierarchical clustering of tumour samples harbouring *TP53* splice mutations. The results revealed that tumours could be broadly categorized into two distinct clusters based on the expression patterns of p53 signalling-related genes (Figure 4A).

**Figure 4:**
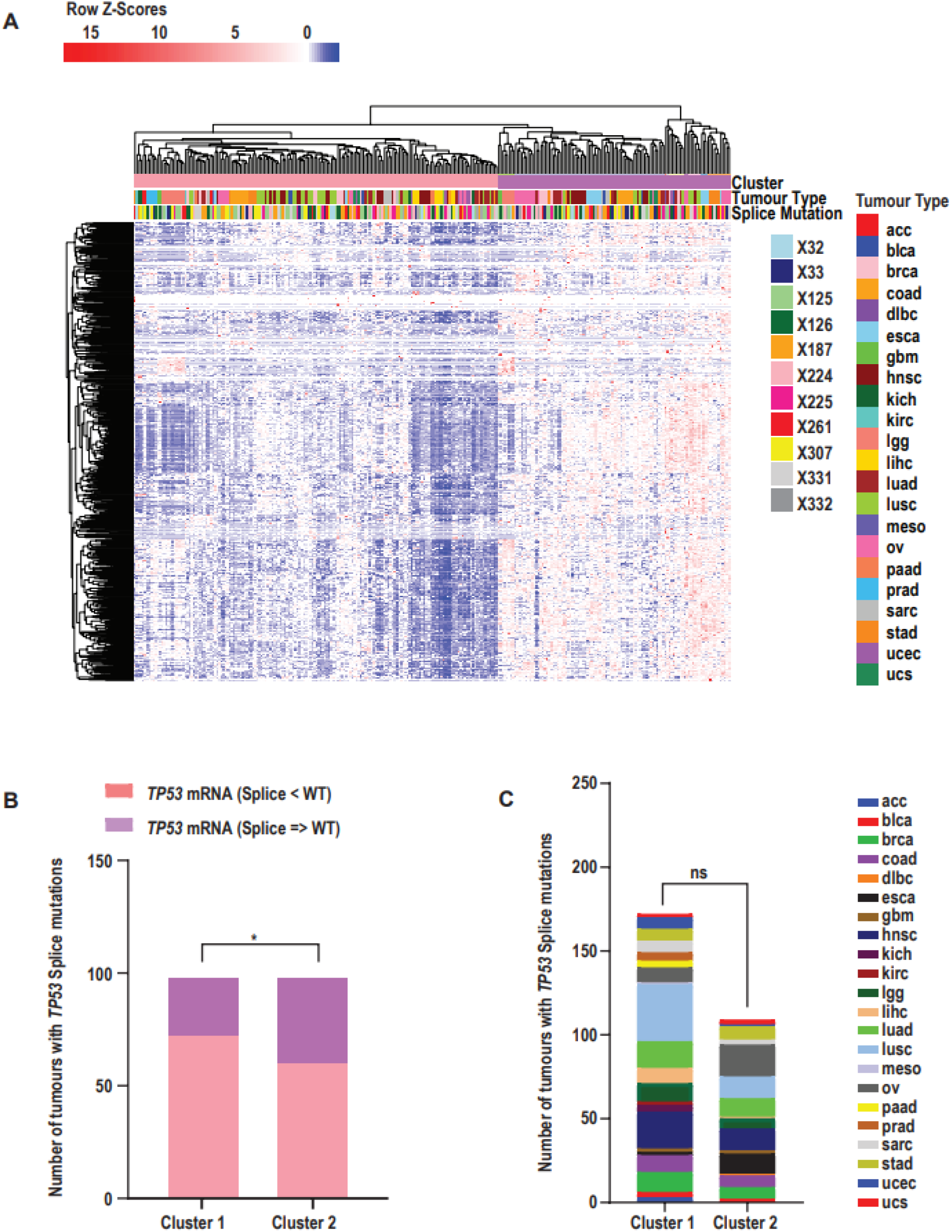
T***P***53 **splice mutations result in aberrant p53 signaling. A.** Pan-cancer heatmap showing mean-centered mRNA expression of genes directly regulated by or associated with p53. Data obtained for the TCGA pan-cancer dataset. Rows represent genes (663); columns represent individual tumour samples (n = 282). Both rows and columns were hierarchically clustered using Euclidean distance and complete linkage. Column sidebars indicate: (i) two expression clusters identified by k-means clustering (Cluster 1: widespread downregulation of p53 signaling genes; Cluster 2: selective downregulation with concurrent upregulation of others), (ii) tumour type, and (iii) presence of TP53 splice mutations. B. Distribution of tumours with TP53 splice mutations based on p53 mRNA levels compared to tumours with WT-TP53. Mutations in X125, X187, X224, X225, X307, and X331 are associated with decreased p53 mRNA expression, while mutations in X32, X33, X126, X261, and X332 show expression levels equal to or greater than WT-TP53. Fisher’s exact test was used to assess statistical significance (p = 0.035; p < 0.05 considered significant). C. Number of tumours with TP53 splice mutations by tumour type in Cluster 1 and Cluster 2. Chi-square test for trend; p = 0.7294 (ns = not significant).

Tumours in Cluster 1 exhibited a global downregulation of genes associated with p53 signalling and a higher proportion of splice mutations for which overall *TP53* mRNA expression was reduced compared to wild- type (WT) *TP53* tumours. In contrast, tumours in Cluster 2 displayed a more heterogeneous expression pattern, characterized by selective downregulation of certain p53 target genes alongside upregulation of others. This cluster also included a higher proportion of splice mutations associated with *TP53* mRNA expression levels that were comparable to or greater than those observed in WT-*TP53* tumours (Figure 4A and 4B). Notably, these clustering patterns and their associated expression profiles were not attributable to tumour anatomical site, indicating that the observed differences in splice mutation frequency and transcriptional impact were not driven by cancer type (Figure 4A and 4C).

These results underscore the transcriptional heterogeneity among *TP53* splice mutations and highlight their potential to differentially disrupt p53 signalling, independent of tumour type.

### Association of individual *TP53* splice mutations with genomic instability

Building on these transcriptional findings, we next assessed whether *TP53* splice mutations are also associated with broader genomic alterations indicative of tumour genomic instability. Specifically, we analysed tumour mutation burden and fraction of genome altered (FGA) using data from the pan-cancer TCGA and MSK cohorts. Tumours with *TP53* mutations exhibited a higher overall mutation count compared to those with WT-*TP53* (Figure 5A). However, mutation burden varied by the type of *TP53* mutation. Tumours with splice site mutations showed significantly higher mutation counts compared to those with missense or nonsense *TP53* mutations. Specific splice mutations, such as X125, were linked to significantly higher mutation counts compared to other splice mutations, except X331 (Figure 5B). Additionally, X331 splice mutations also showed significantly higher mutation counts than X126, 187, 261, and 307 (Figure 5B).

**Figure 5:**
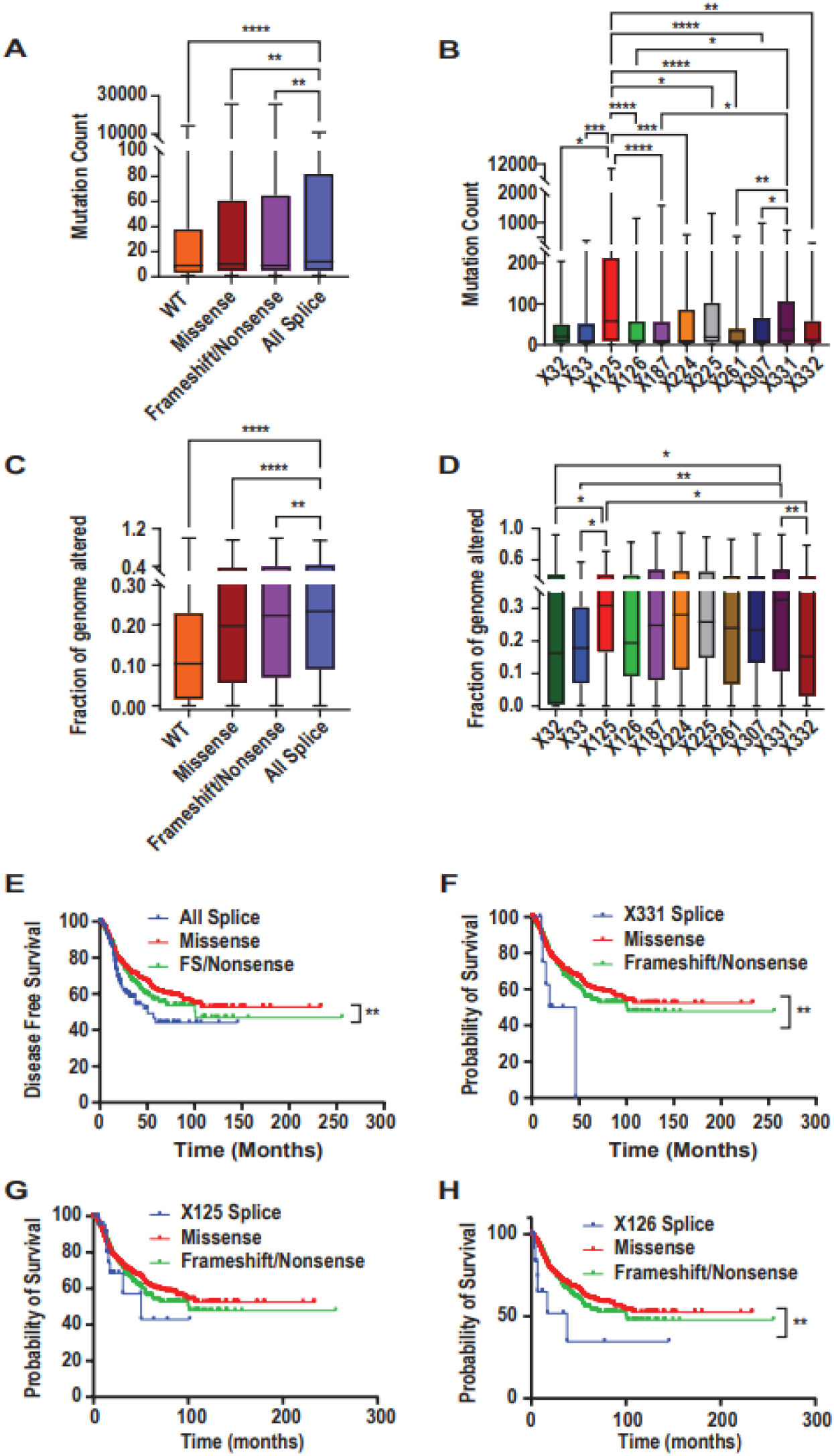
G**e**nomic **instability and disease free survival associated with *TP53* splice mutations. A and C.** Box plot showing the distribution of **A.** Mutation count as a measure of tumour mutation burden (TMB) and **C.** Fraction of the genome altered (FGA), for tumours with either WT-TP53 (n=15075) or containing either, Missense (n=7779), Frameshift/nonsense (n=3115) and Splice (n=835) mutations. B and D. Box plot showing the distribution of B. Mutation count and D. FGA, for tumours with individual TP53 splice mutations including X32 (n=25), X33 (n=30), X125 (n=88), X126 (n=100), X187 (n=134), X224 (n=62), X225 (n=71), X261 (n=78), X307 (n=87), X331 (n=58) and X332 (n=49). A-D. Datasets used TCGA pan-cancer dataset and MSK-MetTropism datasets from cBioPortal. E-H. Kaplan Meier survival plot comparing the disease-free survival of tumors with **E.** All *TP53* splice mutations (n=137), **F.** *TP53* X331 splice mutation (n=9), **G**. TP53 X125 splice mutation (n=24), **H.** *TP53* X126 splice mutation (n=15) against all *TP53* missense (n=1222) and frameshift/nonsense mutations (n=481). Log-rank test was used to determine significance. P < 0.05 was considered significance. **E-H.** Disease free survival was obtained for the TCGA pan-cancer dataset from cBioPortal.

A similar pattern was observed for genomic instability, measured by FGA. Tumours with *TP53* mutations had significantly higher FGA scores than those with WT-*TP53*. Among the mutation types, splice site mutations had the highest FGA scores, surpassing missense, nonsense, or frameshift mutations (Figure 5C). We further analysed the FGA scores across individual splice variants and found that tumours with X331 had the highest median FGA score, followed by X125, X224, X225, X187, X261, X307, X126, X33, X32, and X332 in descending order. Notably, X331 and X125 mutations were linked to significantly higher genomic instability compared to mutations in X32, X33, or X332 (Figure 5D). Tumours with X187, X224, X225, and X307 mutations also had FGA scores comparable to those seen with X125 and X331, suggesting these mutations may also contribute to elevated genomic instability. In contrast, tumours with X126 and X261 mutations had lower FGA scores, though these differences were not statistically significant when compared to X125 and X331 (Figure 5D).

These results reveal that individual *TP53* splice mutations are associated with distinct levels of genomic instability, suggesting potential functional differences that may influence tumour behaviour and patient prognosis.

### Prognostic significance of specific *TP53* splice mutations in cancer

To determine whether these mutation-specific genomic profiles are clinically relevant, we next evaluated the prognostic significance of individual *TP53* splice mutations using disease-free survival data from the pan-cancer TCGA cohort. Tumours with *TP53* mutations had significantly shorter disease-free survival compared to those with WT-*TP53* (Figure S2A). Moreover, tumours with *TP53* splice mutations had even shorter disease-free survival than those with *TP53* missense mutations, but not when compared to *TP53* frameshift or nonsense mutations (Figure 5E). Tumours with X125 and X331 mutations were associated with higher mutation burdens and genomic instability, so we further examined their impact on disease-free survival. Tumours with X331 splice mutations had significantly shorter disease-free survival compared to those with missense, frameshift, or nonsense mutations (Figure 5F). While X125 mutations were linked to higher mutation levels and a higher FGA score, these tumours exhibited similar survival to those with other *TP53* mutation types (Figure 5G). Moreover, X187, X224, X225, X261 and X307 were associated with lower mutation burden compared to X331 and X125 but had similar survival to those with other *TP53* mutation types (Figure S2B). In contrast, tumours with X126 splice mutations, despite having lower mutation burden and FGA scores, were also associated with significantly shorter disease-free survival compared to other *TP53* mutation types (Figure 5H). Importantly, these *TP53* splice mutations were distributed across multiple tumour types, with no single cancer type showing a dominant enrichment (Figure S2C). Therefore, the observed differences in disease-free survival are unlikely to be driven by tumour type–specific effects.

Overall, these findings emphasize the clinical relevance of specific *TP53* splice mutations in predicting disease-free survival, with distinct mutations contributing differently to tumour progression and patient prognosis.

## Discussion

The *TP53* mutation landscape in human tumours is notably diverse and tissue specific. Despite extensive investigation, a substantial knowledge gap remains regarding the biological consequences of *TP53* mutations beyond well-characterized hotspot regions. This limitation poses a significant challenge in leveraging *TP53* status to improve clinical outcomes. Although multiple spliced isoforms of p53 have been identified^1,2,7^, the functional significance of *TP53* splice site mutations is underexplored. This has hindered the clinical application of these variants, despite growing evidence of their impact on tumour biology.

In this study, we performed a comprehensive data mining analysis of *TP53* splice mutations in both somatic and germline tumours. Our results indicate that approximately 7% of all *TP53* mutations are splice mutations, observed across both somatic and germline contexts. Notably, this pattern mirrors that of somatic hotspot missense mutations, which are likewise found in both germline and somatic contexts^27^. Splice mutations were distributed similarly between donor and acceptor sites across various introns, however, in several instances, the base substitution patterns leading to splice mutations differed between germline and somatic tumours. Moreover, germline tumours exhibited a higher frequency of acceptor site mutations driven by transition base substitutions. These observations underscore distinct mutational mechanisms operating in germline versus somatic contexts.

We next evaluated the association of splice mutations with *TP53* mRNA and protein expression, p53 signalling, genomic instability, and clinical prognosis in somatic tumours. *TP53* splice mutations were associated with heterogeneous effects on mRNA and protein levels when compared to missense or frameshift or nonsense mutations^26^. While some mutations led to reduced gene expression, others such as X126 and X261, were linked to increased *TP53* mRNA and protein levels. This upregulation may reflect enhanced expression of specific p53 isoforms^1,2,7^, stabilization of truncated protein variants^28–30^, or alternative splicing events that evade nonsense-mediated decay (NMD) mechanisms^31^. Notably, the X332 mutation demonstrated reduced mRNA expression yet elevated protein levels, suggesting post-transcriptional mechanisms modulating protein stability^28–30^. These findings indicate that *TP53* splice mutations exert complex, context-dependent effects on gene expression, with potential consequences for tumour behaviour through isoform regulation, protein stabilization, and NMD bypass.

Altered p53 expression can disrupt its canonical signalling pathway, influencing transcriptional regulation and tumour progression. Our pan-cancer RNA sequencing analysis of p53 target genes identified two distinct transcriptional clusters. One cluster exhibited global downregulation of p53 targets and was enriched for X125 and X331 mutations, both associated with reduced *TP53* mRNA and protein levels. The second cluster showed selective gene regulation, including both downregulated and upregulated targets, and was enriched for X126 and X261 mutations, both linked to elevated *TP53* expression. These transcriptional profiles appeared independent of tumour tissue of origin, suggesting that the functional consequences of splice mutations are cancer-type agnostic. This variability further emphasizes that *TP53* splice mutations modulate p53 signalling in diverse ways, potentially influencing tumour biology and response to therapy.

As *TP53* is widely recognized as the “guardian of the genome”^32^ and a key suppressor of genomic instability in *in vitro* models, animal studies, and human cancers^33^, we investigated whether splice mutations affect genomic stability. Tumours with *TP53* splice mutations exhibited significantly higher mutation burdens and increased genomic instability, as reflected by elevated FGA scores. X125 and X331 mutations were associated with the highest mutation counts and instability, consistent with their transcriptional suppression of p53^33^. Furthermore, the X331 mutation correlated with significantly poorer overall survival compared to other *TP53* mutation types. Similarly, X125 mutations were associated with poor prognosis, consistent with general *TP53*-deficient phenotypes. These data suggest that certain splice mutations may drive tumour progression by inducing a mutator phenotype, accelerating the accumulation of tumour heterogeneity, oncogenic alterations and worsening clinical outcomes.

Conversely, splice mutations such as X126, which are associated with elevated p53 mRNA and protein levels, can selectively repress the transcription of canonical p53 target genes. This selective loss of transcriptional activity may undermine p53’s tumour suppressive functions while simultaneously enabling gain-of-function effects that promote tumour progression and survival^34^. This is notable because, despite relatively lower genomic instability, X126 mutations were associated with significantly worse survival, indicating that tumour progression can occur independently of genome destabilization in the context of splice-driven p53 isoform changes.

Our findings underscore the clinical relevance of *TP53* splice mutations, despite the limited number of cases identified. This scarcity may reflect historical sampling biases, where many studies utilized exome sequencing panels that excluded noncoding intronic regions. Moreover, earlier datasets often focused solely on the *TP53* DNA-binding domain, potentially underreporting splice site variants. Therefore, the actual frequency and functional diversity of *TP53* splice mutations may be underestimated. Additionally, given the differential impact of individual splice mutations on *TP53* mRNA, protein expression, and isoform composition, further functional validation is needed. Developing *in vitro* and *in vivo* models will be critical for elucidating how specific splice mutations alter the p53 isoform landscape, influence protein stability, and contribute to genomic instability and prognosis.

In summary, *TP53* splice mutations represent a distinct and clinically meaningful class of alterations in cancer. Their variable effects on gene expression and genomic stability highlight the need to consider them separately in both research and clinical contexts. Future studies should aim to uncover the molecular mechanisms underlying these mutations and explore therapeutic strategies tailored to their unique biological impacts.

Figure S1, related to Figure 3

**Figure S1: Impact of *TP53* splice mutations on p53 protein levels. A-B.** Density plot of p53 protein levels from the RPPA data of the TCGA pan-cancer dataset. **A.** Tumors with WT-p53 (n=4634, grey) versus different mutation types (red), left panel: Missense (n=1762), middle panel: frameshift/nonsense (n=702) and right panel: splice (n=205). **B.** Tumors with WT-p53 (n=4634, grey) versus *TP53* splice mutations (red) including: X32 (n=5), X33 (n=13), X125 (n=35), X126 (n=22), X187 (n=32), X224 (n=11), X225 (n=22), X261 (n=15), X307 (n=15), X331 (n=19) and X332 (n=12). Kolmogorov-Smirnov (KS) test was used to determine significance. p < 0.05 was considered significant.

Figure S2, related to Figure 5

**Figure S2: Distribution of *TP53* splice mutation and association with disease free survival. A-B.** Kaplan Meier survival plot comparing the disease-free survival of tumours with **A**. WT-*TP53* mutations (n=3474) versus all *TP53* mutations (n=1864) and **B**. *TP53* X187 (n=20), X224 (n=8), X225 (n=12), X261 (n=10), or X307 (n=13) splice mutations against all *TP53* missense (n=1222) and frameshift/nonsense (n=481) mutations. Log-rank test was used to determine significance. P < 0.05 was considered significance. **A-B.** Disease free survival was obtained for the TCGA pan-cancer dataset from cBioPortal. **C.** Bar plot showing the percentage of individual *TP53* splice mutations found in bladder, breast, colorectal, esophagogastric, head and neck, non-small cell lung cancer, renal cell cancers, melanoma and others.

## Acknowledgements

This work was supported by the Emerging First Researcher grant by the Health Research Council of New Zealand (20/638), Sir Charles Hercus Fellowship by the Health Research Council of New Zealand (HRC 21/030), Health Research Council of New Zealand Program Grant (HRC 23/470) University of Otago Research Funds (ORG 0123-0624), MWC4078.

## Author Contribution

AAB, DS, NFK performed experiments, analysed and assembled the data and contributed to the writing and reviewing the manuscript, AJ, TS and AB assisted with data interpretation and reviewed the manuscript, SM conceptualised and designed the study, interpreted the data, supervised the study, wrote and reviewed the manuscript.

