## Supplementary figures and images for "*In-Silico* Characterization of *TP53* Splice Mutations in Somatic and Germline Tumours"

### Supplementary Figure 1

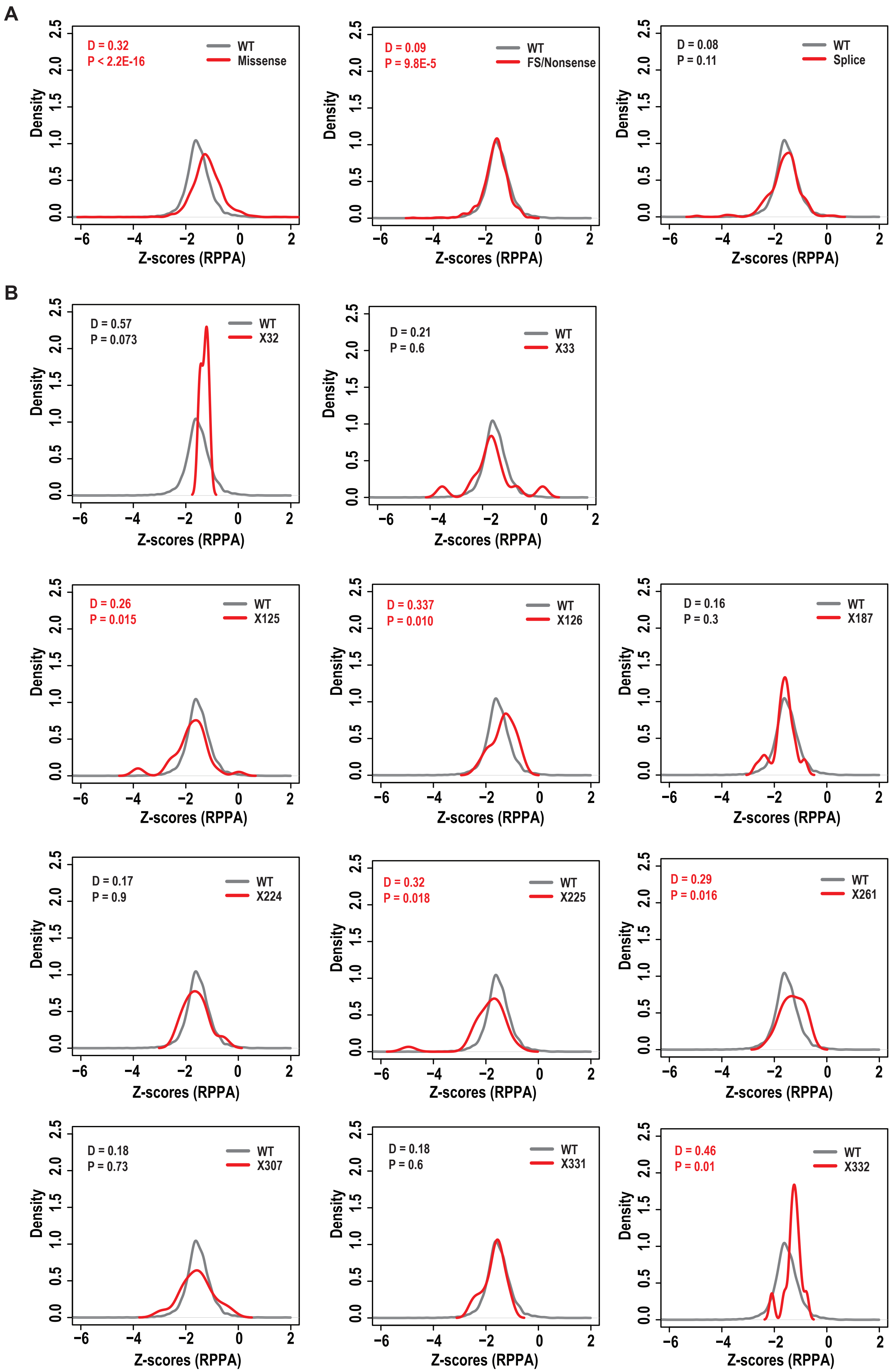

### Supplementary Figure 2

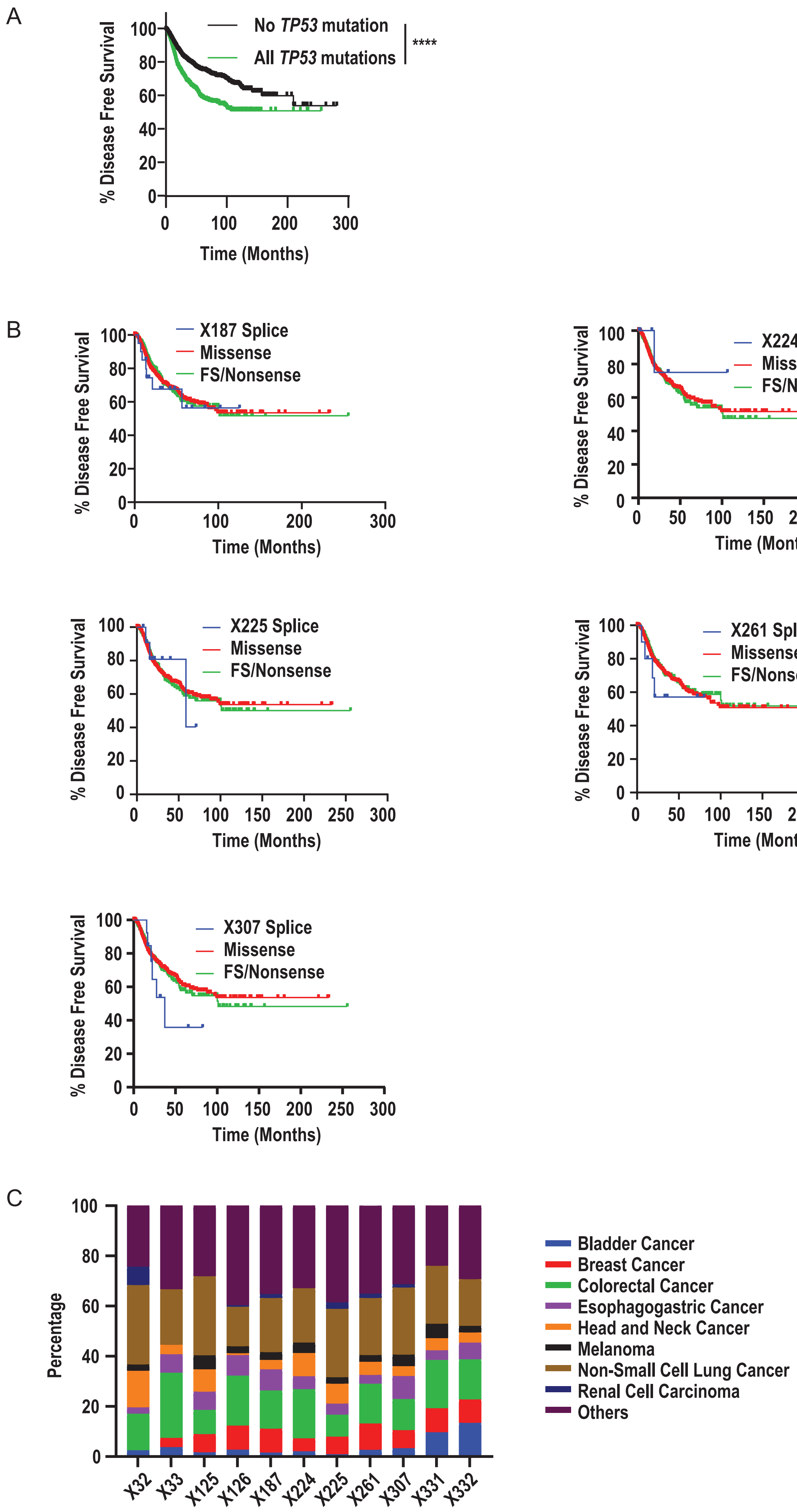
