## Supplementary material for "*In-Silico* Characterization of *TP53* Splice Mutations in Somatic and Germline Tumours": Table 1 and Supplementary Table 1

**Table 1:** Summarises reported *TP53* mutations in splice region, their frequency and type of variant from somatic and germline datasets. Somatic datasets: TCGA Pan-Atlas Cancer Dataset, MSK-MetTropism Metastatic Dataset and Tumour Mutation Burden (TMB) and Immunotherapy Dataset. Germline dataset: IARC database. SNP – single nucleotide polymorphism, DEL – deletion, INS – insertion, AMP – amplification.

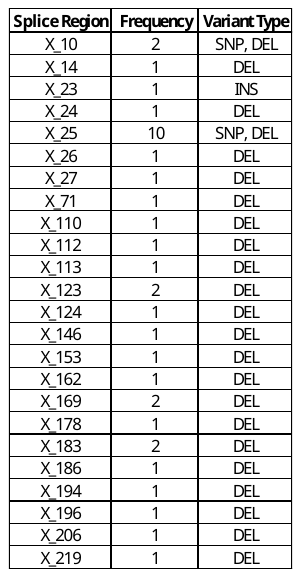

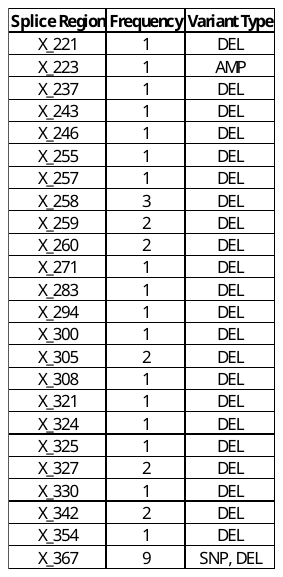

**Table S1. List of *TP53* target genes or those associated with *TP53* expression from ARCHS4 RNA-seq gene-gene co-expression matrix, Enrichr gene-gene co-occurrence matrix, Tagger literature gene-gene co-mentions matrix, and GeneRIF literature gene-gene co-mentions matrix.**

| Combined List | | | | | | | | | | | | | |
| --- | --- | --- | --- | --- | --- | --- | --- | --- | --- | --- | --- | --- | --- |
| AR | DDB2 | KIF20A | RAVER1 | ZDHHC12 | PAGR1 | ZNF202 | ZNF561 | ACTB | CTLA4 | JUN | POLE | ERCC2 | RCHY1 |
| AEN | DDIAS | KIF22 | RCC1 | ZNF217 | PAICSP4 | ZNF212 | ZNF562 | AFP | CTNNB1 | KDR | POU5F1 | ESR2 | RELA |
| ALYREF | DDX12P | KIF23 | RCC2 | ZNF581 | PGBD4 | ZNF225 | ZNF565 | AKT1 | CXCL8 | KEAP1 | PPARG | FAM25C | RPL11 |
| APEX1 | DDX39A | KIF2C | REEP4 | ZWINT | PIPSL | ZNF227 | ZNF566 | ALB | CYCS | KIT | PRKDC | FAS | RPL5 |
| APEX2 | DKC1 | KIFC1 | RFWD3 | ALDH7A1P1 | PMS2P1 | ZNF230 | ZNF567 | ALK | DICER1 | KLK3 | PROM1 | FHIT | S100A4 |
| APRT | DLGAP5 | KPNA2 | RMI2 | AMELY | PMS2P2 | ZNF234 | ZNF574 | ANXA5 | DNMT3A | KRAS | PTEN | GDF15 | S100B |
| ARHGAP11A | DNMT1 | LIG1 | RNASEH2A | CA5BP1 | POT1 | ZNF260 | ZNF576 | ANXA8 | DNTT | KRT19 | PTGS2 | GSTM1 | SAMD4B |
| ARHGEF19 | DPP3 | LMNB1 | RPA1 | CENPBD1 | PSME2P2 | ZNF263 | ZNF584 | APAF1 | E2F1 | KRT20 | PTPRC | GSTP1 | SCO2 |
| ASF1B | DTL | LMNB2 | RPL12 | CHMP4A | RBM23 | ZNF264 | ZNF586 | ARID1A | EGF | KRT5 | PXDN | GSTT1 | SERPINB5 |
| AURKB | DTYMK | LRR1 | RPL18 | CT47A6 | RHEBP1 | ZNF268 | ZNF587 | ASXL1 | EGFR | KRT7 | PXDNL | HIPK2 | SERPINE1 |
| BAX | E2F2 | MCM2 | RPL18A | CYCSP55 | RHEBP2 | ZNF274 | ZNF614 | ATM | EP300 | LIAT1 | RAD50 | HNRNPK | SF3B1 |
| BCL2L12 | EFNA4 | MCM3 | RPL7A | DBIL5P | RHOQP1 | ZNF282 | ZNF616 | ATRX | EPCAM | MAP2K1 | RB1 | HRK | SFN |
| BIRC5 | EIF3D | MCM5 | RPLP0 | DEFB104B | RNPS1P1 | ZNF284 | ZNF619 | B2M | ERBB2 | MAPK3 | RET | HSPA9 | SIRT3 |
| BUB1 | EIF4A1 | MCM6 | RPS16 | DEFB105A | SAA3P | ZNF286A | ZNF620 | BCL2 | ESR1 | MCL1 | RPS6KB1 | IFI16 | SP1 |
| BUB1B | EIF4EBP1 | MCM7 | RPS19 | DEFB105B | SELV | ZNF3 | ZNF623 | BCL2L1 | FASLG | MDM2 | RUNX1 | ING1 | THBS1 |
| BYSL | EIF5A | MELK | RPS2 | DEFB119 | SEPT7P9 | ZNF304 | ZNF625 | BCL2L11 | FBXW7 | MDM4 | SDHC | KAT2B | TIGAR |
| C10orf2 | ESPL1 | MGME1 | RPS3 | DEPDC4 | SMN2 | ZNF317 | ZNF639 | BCL6 | FGF2 | MET | SIRT1 | KAT5 | TNFSF10 |
| C17orf53 | EXO1 | MKI67 | RPS4X | FAM149B1 | SMURF2P1 | ZNF318 | ZNF646 | BECN1 | FGFR3 | MGMT | SLCO6A1 | KLF4 | TP63 |
| C19orf48 | EZH2 | MTA2 | RPSA | FAM200A | SPACA5B | ZNF319 | ZNF668 | BRAF | FLT3 | MLH1 | SMAD4 | KLF5 | TP73 |
| C19orf54 | FAM60A | MTFR2 | RRM2 | GH1 | STX4 | ZNF324 | ZNF669 | BRCA1 | FN1 | MME | SNAI1 | KMT2D | TWIST1 |
| CAD | FAM64A | MYBL2 | RUVBL1 | GH2 | TCEB3B | ZNF324B | ZNF672 | BRCA2 | FOS | MMP2 | SOD2 | KMT5A | UBE3A |
| CASP2 | FAM86C1 | MYC | SF3B4 | GNRHR2 | THAP9 | ZNF330 | ZNF684 | BTK | FOXO3 | MMP9 | SOX2 | LINC00385 | USP7 |
| CBX2 | FANCA | NCAPD2 | SHMT2 | HIGD2B | TIGD6 | ZNF33A | ZNF687 | CASP3 | GADD45A | MMUT | SRC | LINC01761 | VEGFA |
| CCNB1 | FANCD2 | NCAPH | SLC16A13 | HSPD1P1 | TLK2P1 | ZNF341 | ZNF689 | CASP8 | GAPDH | MSH2 | STAT3 | MAPK1 | VHL |
| CCNB2 | FANCE | NDC80 | SLC1A5 | IFNA1 | TP53BP1 | ZNF343 | ZNF691 | CASP9 | GFAP | MSH6 | STK11 | MAPK14 | XPC |
| CCNF | FANCG | NOB1 | SNRPA | INS | TP53BP2 | ZNF346 | ZNF696 | CCK | GLB1 | MTOR | SYP | MAPK8 | XRCC1 |
| CDC20 | FANCI | NONO | SNRPB | KIAA0100 | TTC31 | ZNF354A | ZNF7 | CCNA1 | GPT | MYCN | TCHP | MEG3 |  |
| CDC25C | FBL | NOP2 | TACC3 | KIAA0391 | TXNRD3NB | ZNF384 | ZNF70 | CCND1 | GSK3B | NANOG | TERT | MIR122 |  |
| CDC45 | FBXO5 | NPM3 | TCF19 | KIAA0586 | UBE2D4 | ZNF394 | ZNF701 | CCNL2 | H2AX | NCAM1 | TET2 | MIR125A |  |
| CDC6 | FOXM1 | NRM | TCF3 | KIAA0753 | VDAC1P1 | ZNF398 | ZNF74 | CD19 | H3C12 | NF1 | TGFB1 | MIR34A |  |
| CDCA2 | GATAD2A | NUP62 | TGIF2 | KIAA1143 | VDAC1P3 | ZNF408 | ZNF740 | CD274 | H3C13 | NFE2L2 | TNF | MIR34B |  |
| CDCA4 | GEMIN4 | NUSAP1 | THOC6 | KRT8P12 | VDAC1P6 | ZNF410 | ZNF746 | CD34 | HDAC9 | NFKB1 | WT1 | MIR34C |  |
| CDCA5 | GINS2 | ORC1 | TIMELESS | LRRC37BP1 | XRCC6P2 | ZNF417 | ZNF747 | CD38 | HIF1A | NFKBIA | XIAP | MTHFR |  |
| CDCA7 | GINS4 | PABPC1 | TK1 | MCTS2P | YY1AP1 | ZNF426 | ZNF749 | CD4 | HMOX1 | NOTCH1 | ABCB1 | NBN |  |
| CDCA8 | GRWD1 | PCNA | TNFRSF10B | METTL2A | ZBED1 | ZNF444 | ZNF76 | CD44 | HRAS | NPM1 | ADAM11 | NCL |  |
| CDK2 | GSG2 | PFAS | TONSL | METTL2B | ZNF101 | ZNF45 | ZNF764 | CD68 | HSP90AA1 | NRAS | APC | NEDD8 |  |
| CDK4 | GTSE1 | PFN1 | TOP2A | MORC2 | ZNF131 | ZNF473 | ZNF766 | CD8A | HSP90AB1 | PALB2 | ATF3 | NME1 |  |
| CDT1 | HDAC1 | PLK1 | TPX2 | MRPS31P4 | ZNF134 | ZNF48 | ZNF768 | CDH1 | HSPA4 | PARP1 | ATR | NOS2 |  |
| CENPA | HJURP | POC1A | TRAIP | MTERF1 | ZNF142 | ZNF480 | ZNF776 | CDH2 | IDH1 | PDCD1 | AURKA | NQO1 |  |
| CENPH | HMGA1 | POLA2 | TRIM28 | OR13C8 | ZNF146 | ZNF490 | ZNF777 | CDK1 | IDH2 | PDGFRA | BAK1 | NUMB |  |
| CENPO | HN1L | POLD1 | TROAP | OR2AG1 | ZNF155 | ZNF500 | ZNF784 | CDK6 | IFNG | PECAM1 | BBC3 | OGG1 |  |
| CEP55 | HNRNPA1 | PPM1G | TTK | OR2F2 | ZNF16 | ZNF510 | ZNF785 | CDKN1A | IGF1 | PGR | BID | PDCD5 |  |
| CHAF1A | HNRNPAB | PPRC1 | TUBA1C | OR52J3 | ZNF17 | ZNF511 | ZNF79 | CDKN1B | IGF1R | PIK3C3 | CASP6 | PIN1 |  |
| CHEK2 | HNRNPF | PSME3 | TYMS | OR52W1 | ZNF174 | ZNF526 | ZNF799 | CDKN2A | IL10 | PIK3CA | CCNE1 | PML |  |
| CHST14 | HNRNPUL1 | PTBP1 | UBE2C | OR56A5 | ZNF18 | ZNF527 | ZNF805 | CDKN2B | IL1A | PIK3CB | COP1 | PPM1D |  |
| CKS1B | IMPDH2 | PTTG1 | UBE2I | OR5D18 | ZNF180 | ZNF543 | ZNF830 | CDKN3 | IL1B | PIK3CD | CREBBP | PPP1R13L |  |
| CKS2 | IPO4 | RACK1 | UHRF1 | OR5I1 | ZNF197 | ZNF548 | ZNF839 | CEACAM5 | IL2 | PIK3CG | CYP1A1 | PPP2R2A |  |
| DAXX | KIF11 | RAD51 | WRAP53 | OR8B4 | ZNF2 | ZNF551 | ZSCAN32 | CHEK1 | IL6 | PMAIP1 | CYP2E1 | PRKAA1 |  |
| DCTPP1 | KIF18B | RAD54L | XRCC3 | PAAF1 | ZNF200 | ZNF557 | ABL1 | CREB1 | JAK2 | PMS2 | ERCC1 | PTK2 |  |
